# Dissociable effects of feature expectation on saccades and presaccadic perception

**DOI:** 10.64898/2026.07.05.735520

**Authors:** Luan Zimmermann Bortoluzzi, Gustavo Rohenkohl

## Abstract

During active vision, the brain must coordinate where to move the eyes with predictions about upcoming sensory input. Before each saccade, perception is enhanced at the upcoming fixation location, but whether this enhancement depends on expectations about target features remains unknown. Here, participants prepared a saccade to a cued location while reporting the presence and orientation of a brief visual target that appeared either at the saccade goal or at the opposite location. Feature expectation was manipulated across blocks by varying the probability of the two target orientations. Perceptual sensitivity (*d*′) increased when targets were presented at the saccade goal, consistent with presaccadic enhancement, and was also higher for less expected features. However, these effects were independent: feature probability did not alter the magnitude of presaccadic enhancement. Moreover, presaccadic enhancement increased near saccade onset, whereas the advantage for less expected features weakened as movement onset approached. Saccade latency revealed a contrasting pattern. Visual targets presented at the saccade goal delayed movement initiation. This delay depended on feature probability, with longer latencies for unexpected than for expected features only when saccades were directed towards the target. This location-specific effect persisted after accounting for perceptual report, and the latency cost for unexpected features was reproduced in a follow-up experiment. Together, these findings show that feature probability enhanced sensitivity to unexpected information independently of presaccadic enhancement, while selectively delaying saccade initiation towards targets with unexpected features. This dissociation suggests that feature expectation modulates perception and action through functionally distinct forms of visual processing.

## INTRODUCTION

Active vision requires the brain to solve two problems at once: selecting where the eyes should move next and predicting what visual information will be encountered there. Before each saccade, visual processing is enhanced at the upcoming fixation location, a phenomenon known as presaccadic attention (for a review, see Li et al., 2021). At the same time, perception is modulated by expectations about the features of upcoming sensory inputs (for a review, see de Lange et al., 2018). Yet, it remains unclear whether these two forms of perceptual modulation interact during active visual sampling or whether they influence perception independently.

Since the mid-1990s, several studies have shown that visual attention is biased toward the upcoming fixation location before the eyes move (Deubel and Schneider, 1996; Hoffman and Subramaniam, 1995; Schneider and Deubel, 1995; for a review, see Li et al., 2021). This presaccadic shift of attention (PSA) enhances visual perception at the saccade goal, improving the processing of stimuli presented at that location. Over the years, PSA has been shown to modulate several aspects of visual perception, including contrast sensitivity (Rolfs & Carrasco, 2012), spatial-frequency and orientation tuning (Kroell & Rolfs, 2021; Li et al., 2016; Ohl et al., 2017), spatial resolution (Kwak et al., 2023; Li et al., 2019), and face recognition (Wolfe & Whitney, 2014).

Additionally, electrophysiological studies have shown that target selection and saccade preparation modulate neural activity across multiple stages of visual processing, from primary visual cortex (Supèr et al., 2004), to extrastriate visual areas such as V4 (Bichot et al., 2005; Moore et al., 1998; Zhang et al., 2026; Zhou & Desimone, 2011), and higher-level temporal cortex (Sheinberg & Logothetis, 2001). Importantly, these presaccadic modulations are not purely spatial. In area V4, neurons show enhanced presaccadic responses when saccades are directed toward stimuli in their receptive fields, and this activity can preserve feature-selective information about the upcoming saccade target (Moore, 1999; Moore et al., 1998). These findings suggest that visual representations at the saccade goal are strengthened before the movement is executed.

If presaccadic attention strengthens not only the spatial location of the upcoming fixation but also feature-level representations of the saccade target, then expectations about what is likely to appear there may be particularly important during active vision. Expectations also play a central role in visual perception. By exploiting regularities in the environment, the visual system can anticipate likely sensory events and use this information to guide perceptual inference (for a review, see de Lange et al., 2018). Importantly, feature probability can influence perception in multiple ways: expected stimuli may benefit from prior information (Wyart et al., 2012), whereas unexpected or less predictable stimuli may be prioritized because they carry new information about the environment (Allenmark et al., 2015; Saurels et al., 2020; for a review, see Press et al., 2020).

Together, these findings raise a central question: are expectations about upcoming visual features incorporated into the presaccadic enhancement of perception at the saccade goal? Here, we tested this question by independently manipulating saccade direction and the probability of target features during a presaccadic perceptual task. This design allowed us to ask whether feature probability changes the magnitude of presaccadic enhancement or whether saccade preparation and feature probability influence perception independently. If expectations modulate PSA, the perceptual benefit at the saccade goal should depend on whether the upcoming feature is expected or unexpected.

## RESULTS

To test whether feature-based expectation modulates presaccadic perceptual enhancement, we designed a task combining target-orientation probability with a saccade cue (Figure 1A). Participants maintained central fixation until a cue indicated whether they should execute a saccade to the left or right location. During saccade preparation, a visual target - a Gabor patch oriented -45° or 45° relative to vertical - could appear either at the saccade target location or at the opposite location, with equal probability. At the end of each trial, participants indicated whether a visual target was present or absent and, when present, reported the target orientation as clockwise or counterclockwise relative to vertical. Feature expectation was manipulated in a blocked design. The high-probability orientation appeared on 64% of all trials, whereas the low-probability orientation appeared on 16%. The remaining 20% of trials were target-absent catch trials. To ensure stable feature-based expectations, the orientation–probability mapping changed only once, halfway through each session. Figure 1 summarizes the task sequence and experimental contingencies.

**Figure 1:**
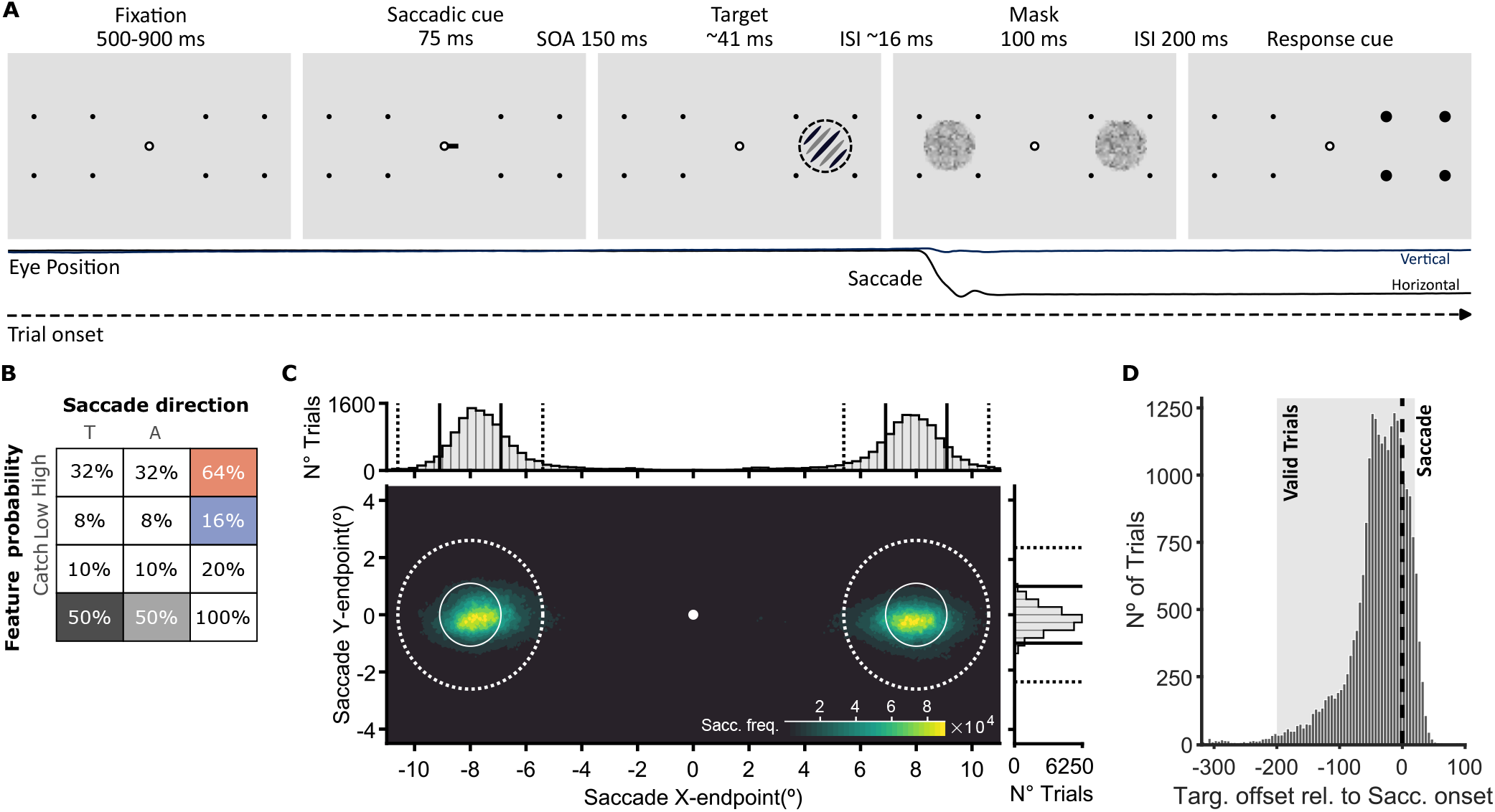
Task design and control parameters. **(A)** Trial example. At the beginning of each trial, participants maintained fixation on the fixation point (FP). After a random period of 500–900 ms, a saccadic cue (black line) was presented for 75 ms, indicating the location to which participants had to execute a saccade (right side in the example). After a stimulus onset asynchrony (SOA) of 150 ms, a visual target (tilted Gabor) was presented for ∼41 ms. On signal-present trials, the target was equally likely to appear on the left or right side of the screen (right side in the example). A visual mask (white noise) was then presented bilaterally after an interstimulus interval (ISI) of ∼ 16 ms, remaining on the screen for 100 ms. In 20% of the trials, no Gabor was presented and only the bilateral masks were shown (*i*.*e*. catch trials). Following an ISI of 200 ms relative to the mask offset, the placeholder dots on one side increased in size to 0.45 dva, serving as a response cue that indicated the side on which participants had to report target presence/absence and, when applicable, target orientation (clockwise or counterclockwise). In this example, the saccadic cue and target locations are congruent. **(B)** Contingency table depicting the proportion (%) of signal-present and signal-absent trials for each experimental condition. Feature expectation was manipulated in a blocked design: the high-probability feature appeared on 64% and the low-probability feature on 16% of the trials. Catch trials were 20% of all trials, split equally across Towards and Away saccade conditions. T and A indicate saccades towards and away from the probed location, respectively. **(C)** Density distribution of saccade endpoints pooled across participants. The y- and x-axes represent vertical and horizontal endpoint positions, respectively. Histograms show the same endpoint data projected onto the horizontal and vertical axes. The solid lines (on both the histogram and density plots) represent the target position on each side, whereas the dashed lines indicate the saccade endpoint-inclusion boundaries. **(D)** Histogram depicting the number of trials as a function of the target offset relative to saccade onset. The vertical dashed black line marks saccade onset. The gray shaded area indicates trials included in the perceptual analyses, for which target offset occurred between -200 and 20 ms relative to saccade onset.

### Presaccadic attention and feature probability independently modulate perceptual sensitivity

Perceptual sensitivity (*d*′) was evaluated for each Saccade Direction condition (Towards and Away from the probed location) and for each Feature Probability (High and Low). Our analysis revealed that perceptual sensitivity increased when participants made perceptual judgments at the saccade-targeted location, relative to the opposite side (Figure 2A, *F* (1, 16) = 29.519, *p* < .001, partial *η*^2^ = .649). There was also a main effect of Feature Probability (Figure 2B, *F* (1, 16) = 44.882, *p* < .001, partial *η*^2^ = .737). Visual sensitivity was higher for the less expected orientations than for the more expected ones. We found no evidence of an interaction between the two factors (Figure 2C, *F* (1, 16) = .6463, *p* = .43, partial *η*^2^ = .039).

**Figure 2:**
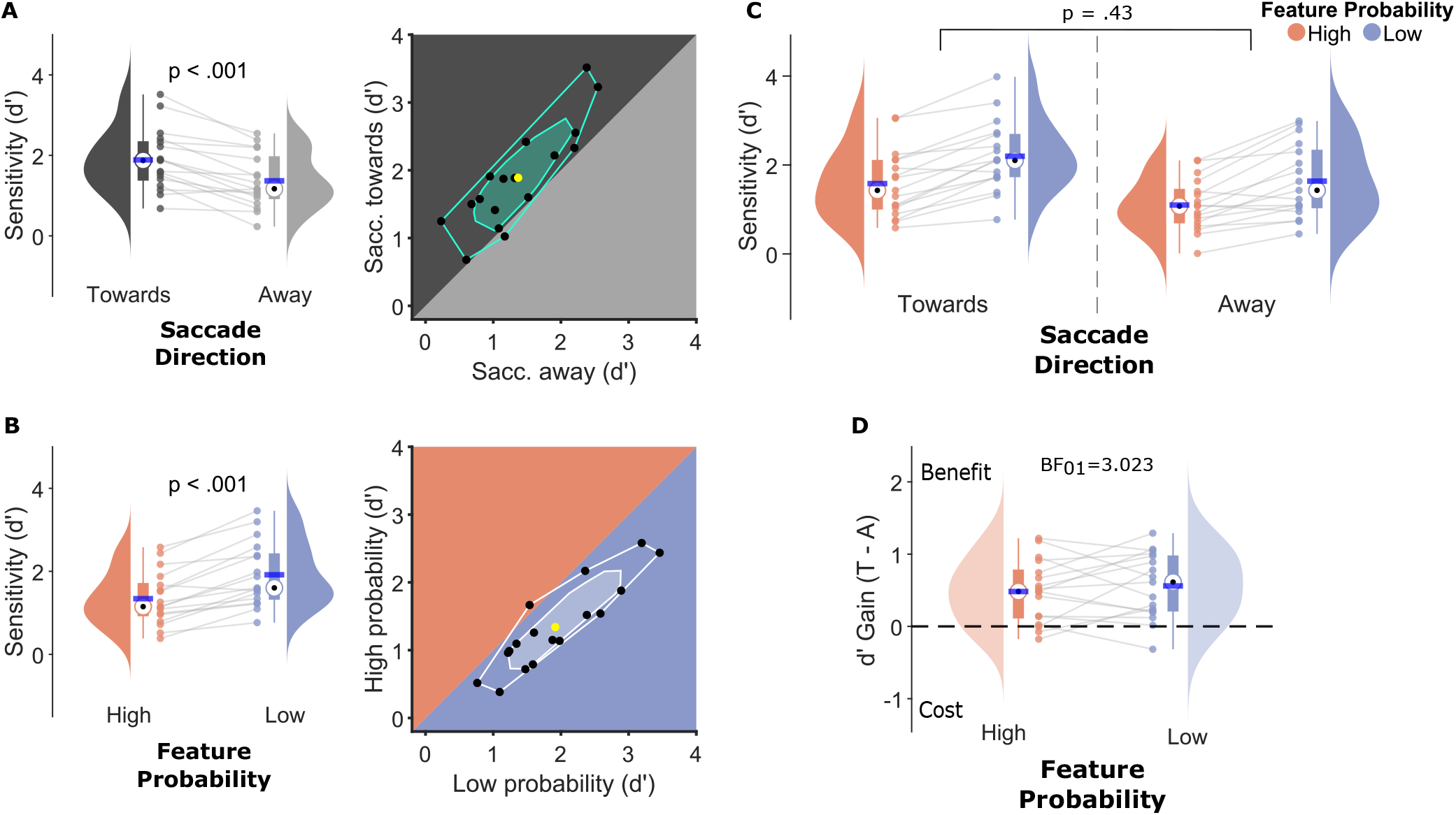
Effects of presaccadic attention and feature expectation on visual sensitivity. **(A)** Perceptual sensitivity (*d*′) as a function of saccade direction relative to the probed location. Towards and Away conditions are shown in dark gray and light gray, respectively. *Left panel:* Half-violin plots depict the distribution of sensitivity values across participants, with individual participants shown as dots. Boxplots show the interquartile range (IQR), whiskers extend to the most extreme values within 1.5 times the IQR, black dots indicate the median, and horizontal blue lines the mean. *Right panel:* Bagplot comparing sensitivity (*d*′) when saccades were directed towards versus away from the probed location. The green polygon (bag) contains 50% of the observations, whereas the surrounding fence contains the remaining non-outlier observations. Black dots indicate individual participants, and the yellow dot indicates the mean. **(B)** Sensitivity (*d*′) as a function of feature probability. High and low feature probability conditions are shown in orange and blue, respectively. *Left panel:* Plotting conventions are the same as in A. *Right panel:* Bagplot comparing sensitivity (*d*′) between high and low feature conditions. **(C)** Sensitivity (*d*′) as a function of feature probability, shown separately for saccades towards and away from the probed location. High and low feature probabilities are shown in orange and blue, respectively. **(D)** Presaccadic gain, computed as the difference in sensitivity between saccades towards and away from the probed location (Towards - Away), for high and low feature probabilities (orange and blue, respectively).

To evaluate the strength of this null effect, we compared the difference in presaccadic gain (Towards - Away) between High and Low Feature Probability conditions using a Bayesian paired t-test. The Bayes Factor indicated moderate evidence in favor of the null hypothesis (Figure 2D, *BF*_01_ = 3.023). The same conditions that increased perceptual sensitivity also produced a more conservative response criterion (*C*), indicating that these effects were accompanied by shifts in response bias (Figure S1). Together, these results suggest that saccade direction and feature probability independently modulate perceptual sensitivity, with higher sensitivity at the saccade target and for less expected features, but no evidence that feature probability affects the magnitude of presaccadic enhancement.

Presaccadic attention begins modulating visual perception before saccade execution and reaches its peak moments before saccade onset (Deubel, 2008). Thus, to ensure that visual sensitivity results were related to presaccadic attention, we only included trials in which target offsets occurred between -200 and 20 ms relative to saccade onsets, as shown in Figure 1D. Nonetheless, even within this latency window, different saccade latency distributions could affect the main analysis. For instance, if the median latency of a given condition was near the target offset (*i*.*e*., 191 ms), visual perception could be more strongly modulated compared to conditions in which the median value was higher. In addition, presaccadic attention is known to be more tightly linked to saccade intention than to saccade execution (Deubel & Schneider, 1996; Wollenberg et al., 2018), and remains present for saccades landing as far as 1.5 dva relative to the target’s edge (Hanning et al., 2019). Considering this, trials were rejected if saccade metrics did not follow these parameters (see Eye Movements in Methods). However, even with this control, differences in saccade endpoint distributions for a given condition could also bias visual sensitivity.

To rule out these possibilities, we ran two separate two-way repeated-measures ANCOVAs with sensitivity as the dependent variable. In one model, we included the slope of the saccade-latency differences across conditions as a covariate; in the other, we included the slope of the saccade-endpoint-error differences across conditions as a covariate (see Data Analysis for details). As in the main analysis, there was a significant main effect of Saccade Direction [(Covariate: Saccade latency; *F* (1, 14) = 50.955, *p* < .001; partial *η*^2^ = .784); (Covariate: Saccade accuracy; (*F* (1, 14) = 25.302, *p* < .001, partial *η*^2^ = .644)] and Feature Probability [(Covariate: Saccade Latency; *F* (1, 14) = 24.513, *p* < .001, partial *η*^2^ = .636); (Covariate: Saccade accuracy; *F* (1, 14) = 11.460, *p* = .004, partial *η*^2^ = .450)] and no significant interaction [(Covariate: Saccade Latency; *F* (1, 14) = 1.007, *p* = .333, partial *η*^2^ = .067); (Covariate: Saccade Accuracy; *F* (1, 14) = .599, *p* = .452, partial *η*^2^ = .041)]. These results indicate that the main sensitivity effects were not explained by condition-specific differences in saccade timing or landing precision.

### Presaccadic enhancement increases whereas feature-probability effects diminish near saccade onset

Previous research has shown that presaccadic visual sensitivity is higher when targets are presented closer to saccade onset (Deubel, 2008; Rolfs & Carrasco, 2012). Although Feature Probability did not interact with Saccade Direction in the main analysis, the effect of feature probability could still depend on when the stimulus was presented relative to saccade onset. To investigate whether these effects changed across the presaccadic interval, we added Presaccadic Timing as a third factor to the main ANOVA, separating trials into Early and Late bins according to whether targets were presented farther from or closer to saccade onset, respectively. The full ANOVA and set of corrected pairwise comparisons are reported in Supplementary Tables 1–3.

First, this analysis revealed that visual sensitivity was higher for targets presented closer to saccade onset (*F* (1, 16) = 43.182, *p* < .001, partial *η*^2^ = .730). This temporal effect was stronger at the saccade-target location, as shown by a significant interaction between Saccade Direction and Presaccadic Timing (*F* (1, 16) = 42.812, *p* < .001, partial *η*^2^ = .728). Post hoc comparisons showed that sensitivity was higher in the Towards-Late than in the Towards-Early condition (*t*(16) = 7.379, *p*_bonf_ *<* .001, *d* = 1.410), indicating that presaccadic enhancement increased closer to saccade onset.

In addition, visual sensitivity was higher in the Towards condition than in the Away condition in the Late time bin (*t*(16) = 7.502, *p*_bonf_ *<* .001, *d* = 1.276, Figure 3A). In the Early time bin, presaccadic enhancement was smaller and did not survive correction for multiple comparisons (*t*(16) = 2.894, *p*_bonf_ = .063, *d* = .344, *BF*_01_ = 1.441).

**Figure 3:**
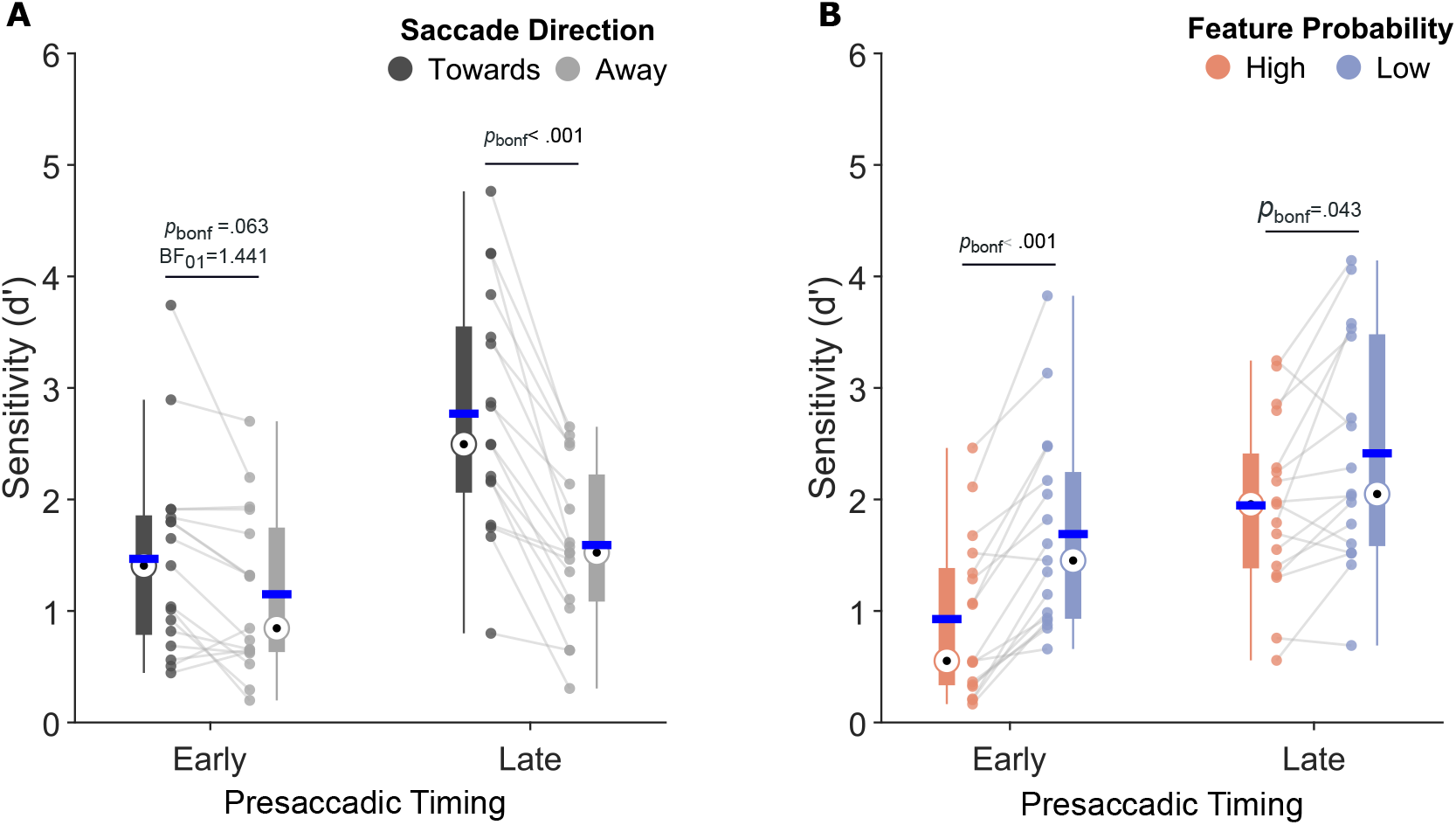
Effects of presaccadic attention and feature expectation on visual sensitivity as a function of presaccadic timing. Presaccadic Timing refers to when the target appeared relative to saccade onset, with Early and Late trials corresponding to targets presented farther from and closer to saccade onset, respectively. **(A)** Sensitivity (*d*′) as a function of Presaccadic Timing, shown separately for saccades towards and away from the probed location. Towards and Away conditions are shown in dark gray and light gray, respectively. Boxplots show the interquartile range (IQR), blue horizontal lines indicate the mean, black dots indicate the median, and whiskers indicate the upper and lower non-outlier limits. *p*_bonf_ values mark the Bonferroni-corrected post hoc comparisons. **(B)** Sensitivity (*d*′) as a function of Presaccadic Timing for high and low feature probability conditions. High and low feature probabilities are shown in orange and blue, respectively. Plotting conventions are the same as in A.

We next examined whether the effect of Feature Probability also changed across the presaccadic interval. In contrast to presaccadic enhancement, which increased closer to saccade onset, the effect of Feature Probability was strongest early in the presaccadic interval and smaller near saccade onset, as shown by a significant interaction between Feature Probability and Presaccadic Timing (*F* (1, 16) = 5.025, *p* = .04, partial *η*^2^ = .239). Post hoc comparisons showed that sensitivity was higher for low-probability than high-probability features in the Early bin (*t*(16) = −6.909, *p*_bonf_ *<* .001, *d* = −.825). In the Late bin, this probability effect was smaller, but still significant (*t*(16) = −3.076, *p*_bonf_ = .043, *d* = −.506, Figure 3B). The three-way interaction between Feature Probability, Saccade Direction, and Presaccadic Timing was not significant (*F* (1,16) = .03, *p* = .865, partial *η*^2^ = .002), indicating that the temporal build-up of presaccadic enhancement did not differ between high and low feature probability conditions. Taken together, these results show that presaccadic enhancement and feature-probability effects followed different temporal profiles. Presaccadic enhancement increased closer to saccade onset, but this increase was not selectively modulated by feature expectation. In contrast, the perceptual advantage for low-probability features was strongest earlier in the presaccadic interval and weakened as saccade onset approached.

### Feature expectation at the saccade location modulates saccade timing

Beyond visual sensitivity, predictions about stimuli presented at the saccade target location may also influence the timing of saccades. So, we examined whether saccade latencies were modulated by target processing and feature probability. Because only signal-present trials contained visual target information, these analyses focused on trials in which a target was presented. In addition, we also included trials in which saccade latencies were not within the presaccadic enhancement period, but that could still be modulated by target processing and feature expectation (See Figure 1D).

Our results showed that saccade latencies were longer when visual targets were presented at the saccade-target location than at the opposite location (*F* (1, 16) = 7.669, *p* = .014, partial *η*^2^ = .324, BF_incl_ = 9.398; Figure 4A), indicating that the presence of visual information at the saccade-target location was associated with delayed saccade initiation. Latencies were also longer for low-probability than high-probability features, as shown by a main effect of Feature Probability (*F* (1, 16) = 4.760, *p* = .044, partial *η*^2^ = .229, BF_incl_ = 2.045). Crucially, feature probability selectively modulated saccade latencies when the visual target appeared at the saccade-target location, as indicated by an interaction between Saccade Direction and Feature Probability (*F* (1, 16) = 5.941, *p* = .027, partial *η*^2^ = .271, BF_incl_ = 5.837). Post hoc comparisons confirmed that low-probability features produced longer latencies than high-probability features when the target appeared at the saccade-target location (*t*(16) = −3.427, *p*_bonf_ = .021, *d* = −.156, *BF*_10_ = 13.09; Figure 4C), but not when it appeared at the opposite location (*t*(16) = −.285, *p*_bonf_ = 1, *d* = .014, *BF*_01_ = 3.873; Figure 4B).

**Figure 4:**
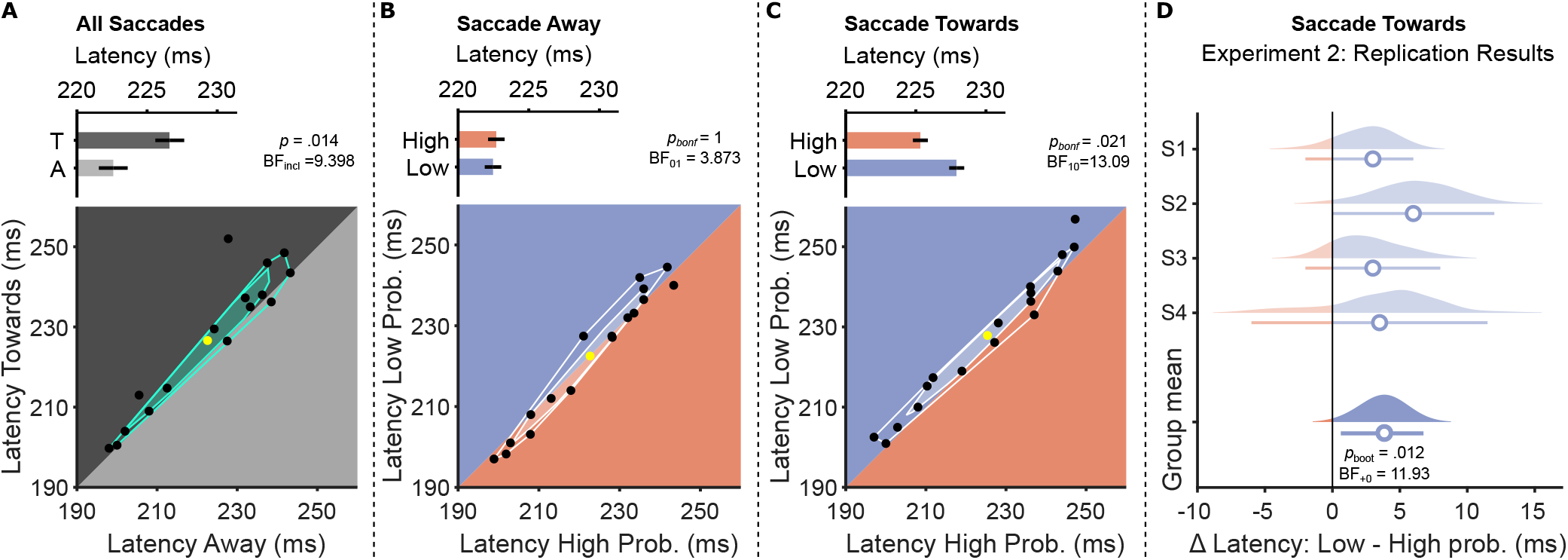
Saccade latencies for signal-present trials. Upper panels show mean saccade latencies (ms), and lower panels show the corresponding bagplots using the same data. Error bars in the upper panels indicate the standard error of the mean (SEM). In the lower panels, black dots indicate individual participants, the polygon (bag) contains 50% of the observations, the surrounding fence contains the remaining non-outlier observations, and the yellow dot indicates the mean. The diagonal line indicates equal saccade latencies between the two conditions. **(A)** Saccade latencies as a function of saccade direction relative to the visual target location. Towards (T) and Away (A) conditions are shown in dark gray and light gray, respectively. **(B)** Saccade latencies for saccades directed away from the visual target location as a function of feature probability. High and low feature probability conditions are shown in orange and blue, respectively. **(C)** Saccade latencies for saccades directed towards the visual target location as a function of feature probability. Paired t-test after outlier exclusion: the difference between towards and away (*p* = .006) and high- and low-probability features for the towards condition (*p* = .004) remained significant, whereas the difference between probabilities in the away condition remained non-significant (*p* = .807). **(D)** Experiment 2: probability effect on saccade latencies at the saccade-target location. Bootstrap distributions for each participant (four in total) and the group mean. Colored clouds represent the within-subject bootstrap resamples (10,000). Blue open circles depict the observed median difference and horizontal colored lines depict the 95% bootstrap confidence intervals (CIs). The vertical black line indicates no latency difference between conditions. Blue and orange colors indicate positive and negative latency effects, respectively, with longer latencies in the low (blue) - and high (orange)-probability conditions. The group mean was evaluated with a within-participant bootstrap (*p*_*boot*_), and *BF*_+0_ shows the directional JZS Bayes factor for the predicted positive effect.

Thus, feature probability modulated saccade initiation selectively for stimuli presented at the saccade-target location, where unexpected features were associated with longer saccade latencies than expected features. We also tested whether other saccade metrics showed similar effects of feature probability. Unlike saccade latency, saccade endpoint dispersion and peak velocity varied with target location, but showed no reliable effects of Feature Probability or Saccade Direction × Feature Probability interactions. These analyses are reported in the Supplementary Text and Supplementary Figure S3.

The results so far show that low-probability features delayed saccade initiation specifically when they appeared at the saccade-target location, whereas no comparable interaction was observed for perceptual sensitivity. This raises the possibility that the oculomotor effect could be partially explained by participants’ perceptual reports. We used a generalized linear mixed model (GLMM) to test whether the location-specific probability effect on saccade latency persisted after accounting for participants’ perceptual reports. The model included Perceptual Report, Saccade Direction, Feature Probability, and the Saccade Direction × Feature Probability interaction as fixed effects, with participant-specific random intercepts and slopes for Perceptual Report, Saccade Direction, and Feature Probability (for details, see Data Analysis).

As predicted, saccade latencies were longer on trials in which perceptual reports were incorrect (*χ*^2^(1) = 32.63, *p* < .001). Critically, the Saccade Direction × Feature Probability interaction remained significant after accounting for Perceptual Report, *χ*^2^(1) = 3.86, *p* = .049, indicating that the location-specific probability effect on saccade latency was not explained by whether participants reported the target correctly. Model-adjusted estimated marginal means from the GLMM showed that, at the saccade-target location, latencies were longer for low-probability features (*M* = 237.6 ms) than high-probability features (*M* = 234.6 ms), high-minus-low estimate = −2.977 ms, *SE* = 1.107, *z* = −2.688, *p*_bonf_ = .014.

In contrast, no reliable probability effect was observed when the target appeared at the opposite location (high probability: *M* = 236.2 ms; low probability: *M* = 236.5 ms), high-minus-low estimate = −0.349 ms, *SE* = 1.144, *z* = −0.305, *p*_bonf_ = 1.000.

These results indicate that ongoing saccade planning is sensitive to expectations about task-relevant visual targets. Given the novelty of this finding, we conducted a second experiment to test whether the latency effect of feature expectation at the saccade-target location could be replicated. The design was similar to the main experiment, but was simplified to focus exclusively on saccade latency and to increase the number of trials per condition. Participants performed a two-alternative forced-choice task in which visual targets were highly visible, with a fixed Michelson contrast of 50% and high mean discrimination accuracy in both probability conditions (High Probability: 92.96%; Low Probability: 92.80%). Additionally, the targets always appeared at the saccade-target location, and no catch trials were included (for details, see Experiment 2 in Methods). Despite these changes, saccade latencies were again longer when the target feature was less probable at the saccade goal. The median latency difference was positive in all four observers. A within-participant bootstrap indicated that the effect was unlikely to arise under a zero or negative latency difference (*p*_boot_ = .012, *d* = 2.70, 95% *CI* = [.62, 6.65]) and a directional JZS Bayes factor provided strong evidence for the predicted positive effect, *BF*_+0_ = 11.93. Thus, a simplified follow-up experiment reproduced the saccade latency cost for less expected target features, indicating that this effect is robust across task variants and does not depend on the perceptual-report demands of the main experiment.

## DISCUSSION

The present findings show that feature prediction and presaccadic attention have distinct consequences for perceptual sensitivity and oculomotor behavior. Consistent with a large body of work on presaccadic attention, perceptual sensitivity was enhanced at locations targeted by saccades (Bortoluzzi et al., 2025; Hanning et al., 2019; Kroell & Rolfs, 2021; Li et al., 2021; Rolfs & Carrasco, 2012). This perceptual enhancement was unaffected by stimulus probability. In contrast, feature probability modulated perceptual sensitivity independently of presaccadic selection. Interestingly, however, unexpected target features increased saccade latency specifically for targets that appeared at the saccade goal. Together, these findings suggest that feature expectations can have distinct effects during active vision, modulating visual sensitivity for perception independently of presaccadic enhancement while delaying action towards unexpected targets.

The absence of an interaction between feature expectation and presaccadic attention should not be interpreted as evidence that presaccadic perceptual enhancement is limited to spatial selection. Neurophysiological and psychophysical studies have consistently shown that presaccadic selection prioritizes feature- and object-level information associated with the selected target (Li et al., 2016; Moore, 1999; Moore et al., 1998). Burrows et al. (2014) showed that selecting a stimulus for a saccade can bias visual processing globally towards the features of the saccade target. Visual-search studies also show that feature-based selection can guide saccadic behavior beyond the immediate saccade endpoint. Zhou and Desimone (2011) found that search-target information was enhanced in FEF and V4 when the target was selected by a subsequent saccade, even though it was not the goal of the immediately upcoming eye movement. Thus, presaccadic priority can reflect not only the location selected for the next eye movement, but also task-relevant object features that guide future saccadic choices. Our findings are not at odds with this feature-rich account of presaccadic selection. Instead, they support the distinction between feature-based target selection and feature prediction (Press et al., 2020; Summerfield & De Lange, 2014; Wyart et al., 2012). From this perspective, presaccadic attention can enhance feature information associated with a selected object, without necessarily drawing on learned expectations about which feature is likely to appear at the saccade target.

Although expectations often facilitate perception of likely events, predictable stimuli can also be perceptually attenuated. This apparent inconsistency has been described as a perceptual prediction paradox: expectations can bias perception towards likely input, but unexpected input may be prioritized when it carries information useful for updating internal models (Press et al., 2020). Our results are consistent with the latter possibility, showing that low-probability stimuli were associated with enhanced perceptual sensitivity (Allenmark et al., 2015; Press et al., 2020; Saurels et al., 2020). Press et al. (2020) have suggested that unexpected inputs in low-intensity signal-detection paradigms may often have low sensory precision and therefore may be less likely to trigger surprise-driven upweighting. Here, however, we found a perceptual facilitation of unexpected stimuli in a low-contrast, masked detection-choice task, suggesting that statistically unexpected visual features can be prioritized even when sensory evidence is weak. Critically, however, this probability effect did not translate into a larger presaccadic benefit at the saccade target.

The time-resolved analysis further clarified this relationship. Presaccadic enhancement increased as targets were presented closer to saccade onset, whereas the perceptual advantage for low-probability features was strongest earlier in the presaccadic interval and became smaller near saccade onset. These divergent temporal profiles raise the possibility that expectation-based and saccade-related influences on perception may be reweighted as movement preparation approaches release. However, this temporal attenuation of the probability effect was not specific to the saccade-target location, and the interaction between feature probability, saccade direction, and presaccadic timing was not significant. This pattern further suggests that feature probability and presaccadic attention contribute separately to perceptual sensitivity, while following distinct temporal dynamics.

Surprisingly, although feature probability did not interact with presaccadic attention in perceptual sensitivity, unexpected features presented at the saccade target delayed saccade initiation. This latency cost was reproduced in a follow-up experiment, suggesting that the effect of feature expectation on saccade timing was robust across task variants and different perceptual demands. This effect can be interpreted in light of accumulator models of saccade latency, although the present task differs from classic probability manipulations in saccade-latency experiments. In the LATER (Linear Approach to Threshold with Ergodic Rate) framework, saccade latency reflects the time required for a decision signal (*S*(*t*)) to rise from an initial level (*S*_0_) to a response threshold (*S*_*T*_). Latency therefore depends on the effective distance to threshold (*θ* = *S*_*T*_ − *S*_0_) and on the rate of rise of the signal (*r*), such that *T ≈ θ/r* (Carpenter & Williams, 1995; Noorani & Carpenter, 2016). In classic LATER studies of saccadic prior probability, expectation is typically manipulated by changing where or whether a peripheral saccade target is likely to appear, with higher prior probability shifting the decision signal closer to threshold and thereby reducing latency (Carpenter, 2004; Carpenter & Williams, 1995; Oswal et al., 2007). Urgency and stimulus information can also alter latency, but through partly distinct model components: urgency can change the response criterion (*θ*) whereas stimulus information is more naturally associated with changes in the rate of rise (*r*) (Noorani & Carpenter, 2016; Reddi & Carpenter, 2000). In the present task, by contrast, the saccade was guided by a central cue, target location was not probabilistically biased, and probability concerned the feature identity of a stimulus appearing while a saccade plan was already being prepared.

Nevertheless, the latency cost for unexpected features can be viewed as a form of oculomotor procrastination: a brief postponement of movement release when newly arriving information at the selected saccade goal violates expectation. In this sense, one interpretation of our results is that learned feature expectations biased the decision process associated with the prepared saccade, such that an unexpected feature at the saccade goal transiently updated or interrupted the evolving decision signal (*S*(*t*)) before movement release. Alternatively, the latency cost may reflect a later effect on motor preparation itself, such as a reduction in the rate of rise of the decision signal (*r*), or an increase in the effective response criterion (*θ*). The fact that feature probability modulated saccade latency without modulating presaccadic perceptual enhancement is more consistent with a motor-decision account of the latency effect than with a simple change in perceptual gain at the saccade target. This interpretation should remain cautious, however, because the present data were not fitted with a LATER model and therefore cannot distinguish between the parameters that could account for the latency effects. Future studies could test this account more directly by applying LATER modeling to similar designs, asking whether feature probability is best captured by a feature-dependent update of the evolving decision signal (*S*(*t*)), by changes in the accumulation rate (*r*), or by changes in the distance to threshold (*θ*).

The finding that feature probability modulated perceptual sensitivity independently of presaccadic enhancement, while selectively delaying saccades towards unexpected target features, can be interpreted within the dual-systems model of vision proposed by Goodale and Milner (Goodale & Milner, 1992). According to this model, visual perception and visually guided action depend on partially distinct computations associated with ventral and dorsal visual streams, respectively (Goodale, 2011; Goodale & Milner, 1992, 2004; Milner & Goodale, 1995, 2008). More specifically, the ventral stream supports object recognition and conscious visual experience, whereas the dorsal stream transforms visual information into coordinates required for action control (Goodale & Milner, 1992; Milner & Goodale, 1995, 2008). Within a dual-systems interpretation, this pattern suggests that predictions about the feature identity of the upcoming saccade target may be incorporated more strongly into visuomotor control than into the perceptual gain associated with presaccadic attention. In other words, feature probability may modulate a dorsal, action-oriented readout of the selected target—reflected in the timing of motor commitment—without modulating the ventral/perceptual readout indexed by presaccadic sensitivity enhancement. Although the present study cannot identify the neural pathway underlying this effect, one possible route is the retinocollicular pathway through the superior colliculus and pulvinar, which has been proposed to provide rapid visual input to oculomotor-control circuits and to support dissociations between perceptual report and eye movements (Spering & Carrasco, 2015). In other words, the present results suggest that prediction is expressed differently across vision-for-perception and vision-for-action. To directly test this hypothesis, future studies could investigate, for example, whether the saccade-latency effect varies with the visibility of the target presented at the saccade goal.

In conclusion, the present study shows that feature expectations and presaccadic attention shape perception and oculomotor behavior, but not through a single unitary mechanism. Presaccadic attention enhanced perceptual sensitivity at the saccade target, and this enhancement became stronger closer to movement onset. Feature probability also modulated perceptual sensitivity, with less expected stimuli producing higher sensitivity. However, learned feature probability did not modulate the magnitude or temporal growth of presaccadic enhancement. Thus, expectation did not simply scale the perceptual priority conferred by preparing a saccade. Instead, feature probability and presaccadic attention contributed separately to visual perception. In contrast, feature probability selectively influenced oculomotor behavior, delaying saccade initiation when unexpected features appeared at the saccade goal, even after accounting for perceptual report and in a seperate experiment. These findings suggest that unexpected features can enhance perceptual sensitivity while simultaneously delaying action towards the saccade target. Prediction during saccade preparation therefore appears to be expressed differently across perceptual and motor readouts: it contributes independently to visual sensitivity, but interacts with saccade preparation in the timing of oculomotor action.

## MATERIAL AND METHODS

### Participants

The experiment was conducted on 19 participants (11 females and 8 males; mean age 26). Except for one author (L.Z.B.), all participants were naive to the purposes of the study. All participants had normal or corrected-to-normal vision, with 14 having dominance in the right eye and 15 being right-handed. All participants gave their informed written consent before participating in the study. The procedures were approved by the local Research Ethics Committee of the University of São Paulo (approval number: 25333219.5.0000.5464), in accordance with the Declaration of Helsinki.

### Apparatus

The stimuli were presented on a high-resolution screen (1920 × 1080, refresh rate 120 Hz, VIEWPixx/3D, VPixx Technologies, Canada) and controlled using the Psychtoolbox toolbox (Kleiner et al., 2007) in MATLAB (MathWorks, 2022). Eye position was monitored at 1 kHz sampling rate using the EyeLink Plus 1000 desktop mount (SR Research) (SR Research Ltd., 2013). Manual responses were recorded using a response box (ResponsePixx).

### Stimuli and Task

At the beginning of each trial, participants were instructed to fixate their eyes on a fixation point (FP) presented on a gray background. The FP consisted of a white dot with a diameter of 0.3 degrees of visual angle (dva), within a black circle with a diameter of 0.6 dva. After 500 ms of successful central fixation (within a 4 dva window), placeholders centered 8 dva from the FP were presented on the left and right sides. Each placeholder was composed of four black dots (each with 0.3 dva in diameter) arranged in a square with a side length of 2.83 dva. The placeholders remained on the screen until the end of the trial. Then, after a random interval of 500-900 ms, a non-predictive (*i*.*e*., 50% validity) cue was presented for 75 ms. The cue was a central black line measuring 0.7 dva in length and 0.15 dva in width and pointed to the left or right side of the screen, indicating the side to which participants were required to execute a saccade. Participants were instructed to make saccades to the center of the placeholder as quickly and accurately as possible. Following a stimulus onset asynchrony (SOA) of 150 ms, a visual target could be presented at the probed or unprobed location for *∼*41 ms. Gabor patches (2.5 cycles per degree; Gaussian envelope of *σ* = 1.1 dva) tilted 45° or - 45° relative to vertical were used as targets. After an interstimulus interval (ISI) of *∼*16 ms relative to the target offset, noise patches (masks) were presented at the center of both placeholders for 100 ms (Gaussian envelope of *σ* = 1.1 dva; pixel noise width of 0.22 dva). Finally, a response cue was presented after an ISI of 200 ms relative to mask offset, indicating the location where participants should give a non-speeded manual response. The response cue was an increase in the size of the placeholder dots (from 0.3 to 0.45 dva). Participants were instructed to give a target-present/target-absent response, and if they reported that the target was presented, they also provided an orientation discrimination response (clockwise or counterclockwise relative to vertical).

### Procedures

The experiment was carried out in a quiet, dimly lit room, with participants sitting comfortably in front of a computer screen with their heads positioned on a chin-rest. The participants performed two experimental sessions of 720 trials, with approximately one hour and thirty minutes of duration. At the beginning of each session, participants were informed that one target orientation was more likely to be presented (64% vs. 16%), irrespective of its spatial location. In 20% of the trials, no target was presented (catch trials, see Figure 1B). Each session consisted of 24 short blocks of 30 trials, and the orientation-probability mapping was shown at the beginning of each block to reinforce the probability of target orientation. To strengthen any feature expectation effect, the probability pattern was switched once halfway through each session. In the second session, the orientation-probability mapping was inverted relative to the first session. This latter manipulation was done in order to control for possible learning effects. Additionally, the orientation-probability mapping was counterbalanced across participants. Participants were instructed to respond using their dominant hand. The top button in the response box was used for target-present response, and the bottom button for target-absent response. Clockwise and counterclockwise discrimination responses were made using the right and left buttons in the response box, and the response mapping was counterbalanced across participants. Finally, to encourage participants to execute the saccade as quickly as possible, the FP’s white dot would always turn orange for saccade latencies higher than 300 ms, or if no saccade was detected. Participants were allowed to rest between blocks.

### Task Training

Before the first experimental session, participants received detailed instructions on how to perform the task. Then, participants performed a first training task, which was identical to the staircase task, except that the stimuli had longer durations (see the Staircase Procedure below). After that, a second training task was run with the same stimulus duration as in the staircase, and participants had to complete a total of 60 trials. For both fixation training tasks, Gabor contrast was adjusted based on the participant’s response. In addition to the fixation violation feedback, participants received another visual feedback regarding the target’s presence, in which the FP’s white dot would turn yellow or blue for target present or absent, respectively. In the second experimental session, only the second training task was executed.

Following fixation training and staircase procedure, the central cue was introduced, and participants completed two short blocks in which they were trained to make quick and accurate saccades. The target-orientation probability manipulation was also introduced at this stage, and visual feedback regarding the presence or absence of the target was provided.

### Staircase Procedure

Before each experimental session, the target contrast was estimated using a 1-up/1-down staircase procedure (Levitt, 1971) using the Palamedes toolbox (Prins & Kingdom, 2018), yielding a detection performance of 50%. The staircase was composed of 60 trials, 20% of which were catch trials. The target contrast was estimated using only the signal-present trials (48 in total). For the first session, the initial Michelson contrast was set at 30% and steps of 5% were used for the correct (hits) and incorrect (misses) responses. For the second session, the initial target contrast was the same as the estimated mean value from the first session’s staircase, and the up-down step size was of 2.5%. Participants repeated the staircase task if their false alarm (FA) rate exceeded 25%. The target contrast used in the main task was the mean value calculated from all trials between the fifth-to-last reversal to the end of the staircase procedure.

The staircase task was very similar to the main task, with some important modifications. First, participants were instructed to maintain fixation on the FP throughout the trial. Second, each target orientation had the same probability of appearance (*i*.*e*., 50%). Lastly, at the end of each trial, the central white dot turned orange if the participant’s eyes deviated from the FP. As in the main task, participants had to report the presence or absence of the target and its orientation. However, for the estimation of contrast thresholds, only the first response (signal-present/signal-absent) was considered. This procedure was implemented based on a pilot experiment that showed that detection and discrimination performance for targets with equal orientation probability was highly correlated. This pattern was also reported elsewhere (see supplemental results in Kroell and Rolfs, 2022).

### 2-ADC model implementation

Perceptual performance was computed based on the mADC model (Sridharan et al., 2014), which gives the visual sensitivity index (*d*′) for each condition and the values of the decision criterion (C). The m-alternative detection choice model was developed specifically for the analysis of behavioral data involving two or more alternative choices in detection tasks. It has been extensively tested and validated in previous studies in humans and non-human primates (Gupta & Sridharan, 2024; Lovejoy & Krauzlis, 2017; Sagar et al., 2019; Sreenivasan & Sridharan, 2019; Sridharan et al., 2017). A detailed description of the model can be found in Sridharan et al., 2014, 2017. Here, we briefly describe how the two-alternative detection choice (2-ADC) model was used to compute sensitivity (*d*′) in this study. Sensitivity (*d*′) is defined as:

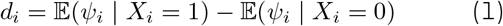

where *ψ*_*i*_ is a bivariate decision variable with its *i*^th^ component representing the sensory evidence for a high-probability feature (*ψ*_1_) and low-probability feature (*ψ*_2_). *X*_*i*_ represents the stimulus event: *X*_*i*_ = 1 denotes sensory signal present (target) with feature probability *i* whereas *X*_*i*_ = 0 denotes sensory signal absent (catch trial). Thus, *d*′ determines how much overlap there is between the signal distribution (*X*_*i*_ = 1) for a given feature (*X*_1_ or *X*_2_) and the noise distribution (*X*_1_ = 0). Conceptually, *d*′ reflects how well participants are able to dissociate signal from noise. The smaller the overlap between the distributions, the larger *d*′ will be. Throughout the text, the terms sensitivity, visual sensitivity, perceptual sensitivity, and *d*′ are used interchangeably.

### Data Analysis

In addition to perceptual sensitivity and decision criterion, we also considered saccade accuracy, saccade latency, and peak eye velocity as dependent variables. Two-way repeated-measures ANOVAs were performed on all dependent variables with Saccade Direction (Towards and Away from the probed location) and Feature Probability (High and Low) as factors. Additionally, to investigate effects as a function of saccade execution time, we performed a median split on saccade latency distributions separately for each participant and each condition. Presaccadic Timing, with Early and Late conditions relative to targets presented farther from or closer to saccade onset, respectively, was then included as a factor in a three-way ANOVA for visual sensitivity.

Normality was tested using the Kolmogorov-Smirnov test, and sphericity violations were verified using Mauchly’s test. Greenhouse-Geisser correction was applied to the p-values when necessary. The effect sizes for the main ANOVAs and post hoc tests were calculated using the partial eta squared (partial *η*^2^) and model-based Cohen’s *d* for paired comparisons, respectively. Bonferroni correction was applied to post hoc analyses.

Bayesian statistics (Peter Rosenfeld & Olson, 2021) were used *ad hoc* to quantify the evidence in favor or against a result. All Bayes Factors (BF) were calculated using a standard Cauchy prior (s = 0.707) for the alternative hypothesis. For Bayesian t-tests, we report BF_01_ and BF_10_ to indicate how much evidence there is in favor of the null or alternative model, respectively. For Bayesian ANOVAs, BF_incl_ and BF_excl_ are reported, indicating how much evidence there is in favor of adding or excluding a given effect (main factor or interaction) from the model.

We also implemented two-way repeated-measures ANCOVAs for visual sensitivity to control for possible effects of saccade latency and saccade endpoint error. For this purpose, we estimated participant-level regression slopes describing differences in saccade latency and endpoint error across the Saccade Direction conditions (Towards and Away from the probed location) and Feature Probability conditions (High and Low). These slope values were then used as covariates in separate ANCOVAs: one for saccade latency and one for saccade endpoint error (see Figure S2).

Outlier detection was performed for visual sensitivity analysis using the Interquartile Range (IQR) rule, in which participants were considered outliers if any of their data points fell outside the lower or higher limits of the IQR. In addition, we also excluded participants who had more than 50% of trials excluded under any of the four conditions for target-present and target-absent trials. In total, two participants were excluded from the main analyses: one by the IQR rule and another for having more than 50% of trials excluded in both target-present and target-absent trial conditions.

To test whether the location-specific effect of feature probability on saccade latency could be explained by participants’ perceptual reports, we fitted a generalized linear mixed model (GLMM) to trial-level saccade latencies. Because saccade latencies were positively skewed, we first evaluated the inverse-Gaussian distribution by fitting inverse-Gaussian distributions separately to each participant’s latency distribution. These fits provided a reasonable approximation to the observed distributions (Kolmogorov–Smirnov statistic: mean = .096, range = .054–.134), consistent with the use of an inverse-Gaussian response distribution in the GLMM (Figure S4). The GLMM used an inverse-Gaussian response distribution with an identity link and included Saccade Direction, Feature Probability, their interaction, and Perceptual Report as fixed effects. Saccade Direction indicated whether the visual target appeared at the saccade-target location or at the opposite location, Feature Probability indicated whether the target feature was of high or low probability, and Perceptual Report indicated whether the trial was correct or incorrect. Trials were classified as correct when participants reported the stimulus and made the correct discrimination, and as incorrect when participants either reported no stimulus or made an incorrect discrimination. The model included participant-specific random intercepts and random slopes for Saccade Direction, Feature Probability, and Perceptual Report, with correlations among random effects estimated. Categorical predictors were sum-coded, fixed effects were evaluated using Type III likelihood-ratio tests, and follow-up comparisons were performed on model-adjusted estimated marginal means.

Statistical data analyses were performed on MATLAB (R2022b) (MathWorks, 2022) and JASP (Version 0.19.3) [Computer software] (JASP Team, 2025).

### Eye Movements

Eye movement data were analyzed using a velocity-based saccade detection algorithm (Engbert & Mergenthaler, 2006). Eye velocities were computed using a moving average procedure, and saccades were detected when the median velocity was exceeded by 5 SDs for at least 8 ms. Saccade onsets between 170 and 190 ms were only considered if saccade offsets were equal to or above 200 ms, ensuring that participants did not receive any target information upon saccade landing, since target offset was at *∼*191 ms from cue onset. Trials were excluded if saccade latencies were above 390 ms or below 170 ms, or if no saccade was detected. We also excluded trials in which participants did not execute a saccade to the cued side and if the saccade endpoint fell 1.5 dva outside the target border (Hanning et al., 2019). In total, 1,945 trials (7.95%) were excluded out of 24,480 trials from the 17 participants.

### Experiment 2

We conducted an additional experiment to assess whether the effects of feature expectation on saccade latencies at the target location could be replicated. To do so, we implemented a two-alternative forced choice (2AFC) discrimination task that was similar to that of Experiment 1 (see Figure 1A and Stimuli and Task), except for the following modifications: The visual target was always presented at the cued location, either at the left or right sides of the FP with equal probability across trials; the visual target was presented in every trial (*i*.*e*., no catch trials) with a fixed 50% Michelson contrast. Four participants from Experiment 1 -including one author (L.Z.B.)-performed this task. Each participant completed a total of 720 trials. In total, 243 trials (8.23%) were excluded out of 2,880 trials from the four participants (for details on trial exclusion criteria, see Eye Movements)).

For statistical inference, a nonparametric bootstrap analysis comparing low- and high-probability conditions was conducted. First, trials were separated into high- and low-probability conditions for each participant. We then estimated the median latency in each condition and computed the latency difference as Low minus High probability. To estimate the latency-difference distribution, the latency values were resampled (with replacement) separately for each condition and participant. This procedure was repeated 10,000 times, creating a latency-difference distribution for each participant (see S1–S4 density clouds in Figure 4D). At the group level, the bootstrap distribution was obtained by averaging the participant-level latency differences on each bootstrap iteration (see Group mean density cloud in Figure 4D). To estimate the statistical significance of the latency-difference effect, a one-tailed p-value (*p*_boot_) was computed based on the proportion of bootstrap iterations at the group level in which the latency difference (Low minus High) was lower or equal to zero. The effect size (Cohen’s *d*) was calculated based on the mean of the group latency-difference distribution divided by its standard deviation. In addition to the bootstrap analysis, we quantified evidence for the predicted positive effect using a directional Jeffreys–Zellner–Siow (JZS) Bayes factor (*BF*_+0_). The Bayes factor was computed from the participant-level latency differences using a one-sample Bayesian *t*-test with a default Cauchy prior on the standardized effect size 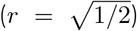. The alternative hypothesis was restricted to the predicted direction, corresponding to longer latencies in the low-probability than in the high-probability condition.

## ACKNOWLEDGMENTS

The authors thank Gabriela Mueller de Melo and João Gabriel Borges da Silva for their helpful comments on the manuscript and lab colleagues for the discussions throughout the development of this study. Supported by IDOR/Pioneer Science Initiative and Conselho Nacional de Desenvolvimento Científico e Tecnológico (CNPq) - grant 140084/2023-1.

## AUTHOR CONTRIBUTIONS

Conceptualization and methodology: L.Z.B., G.R.

Research: L.Z.B., G.R. Software: L.Z.B.

Data collection: L.Z.B.

Data analysis: L.Z.B., G.R.

Visualization: L.Z.B., G.R.

Writing: L.Z.B., G.R.

Manuscript Review: L.Z.B., G.R.

Funding acquisition: L.Z.B., G.R.

## DATA AVAILABILITY

The datasets generated in this study are available from the corresponding author upon reasonable request.

## AUTHOR COMPETING INTERESTS

The authors declare no conflict of interest.

## Notes

### Competing Interest Statement

The authors have declared no competing interest.

