## Supplemental text for "Dissociable effects of feature expectation on saccades and presaccadic perception"

### SUPPLEMENTARY INFORMATION

#### Presaccadic attention and unexpected stimuli lead to more conservative responses

In Signal Detection Theory (SDT) experiments, the Decision Criterion (C) is defined as the observer's bias to decide between signal present vs signal absent, considering the possible benefits or costs when correctly reporting the signal presence (hits) or wrongly reporting its presence when no signal is presented (false alarms). A conservative response (C > 0) indicates that the subject needs more evidence to report the presence of the target, while a liberal response (C < 0) indicates that less evidence is needed.

In the present study, subjects had to give one or two perceptual responses in the end of each trial. The first was a signal-present/signal-absent response, typical of detection experiments. If subjects reported the signal presence, a second response should be given related to the target's tilt (clockwise or counterclockwise), typical of discrimination experiments. Subjects were encouraged to report the signal presence if they were sure of the target appearance, even when they did not know its orientation or were not sure. Interestingly, the majority of the subjects were more conservative in their responses, as shown in Figure 1. In addition to this overall conservative bias, a two-way repeated-measures ANOVA showed that presaccadic attention led to more conservative responses (Figure 1A, $F(1,16) = 19.535$, $p < .001$, partial $\eta^2 = .5485$). There was also a significant main effect of target probability (Figure 1B, $F(1,16) = 93.594$, $p < .001$, partial $\eta^2 = .854$), with a more conservative bias towards less expected features. There was no interaction between factors (Figure 1C, $F(1,16) = .020$, $p = .8882$, partial $\eta^2 = .0013$).

#### Target appearance modulates saccade endpoints and peak eye velocity for signal-present trials

We were also interested in investigating the effects of target appearance and feature probability on saccade endpoint distribution and peak eye velocity. First, we performed a two-way repeated-measures ANOVA with saccade endpoint error as a dependent variable. Endpoint error was calculated by the distance of the saccade endpoint to the center of the target and mask locations. There was no significant effect of Feature Probability ($F(1,16) = .0197$, $p = .89$, partial $\eta^2 = .0012$) nor an interaction between Feature Probability and Saccade Direction ($F(1,16) = 3.1958$, $p = .0927$, partial $\eta^2 = .1665$). Interestingly, however, there was a significant effect of Saccade Direction, where endpoints errors were significantly lower in the Towards condition than in the Away condition (Figure 3A; $F(1,16) = 20.963$, $p = .0003$, partial $\eta^2 = .5671$).

Lastly, we investigated if saccade direction and feature probability might have modulated peak eye velocities (°/s) in signal-present trials. A two-way repeated-measures ANOVA with peak eye velocity as dependent variable did not reveal a significant effect of Feature Probability ($F(1,16) = .0042$, $p = .9487$, partial $\eta^2 = .0003$) nor an interaction between the factors ($F = (1,16) = 2.4808$, $p = .1348$, partial $\eta^2 = .1342$). Interestingly, there was a significant effect of Saccade Direction, with peak eye velocity being higher in the towards condition (Figure 3B; $F(1,16) = 9.714$, $p = .0066$, partial $\eta^2 =$

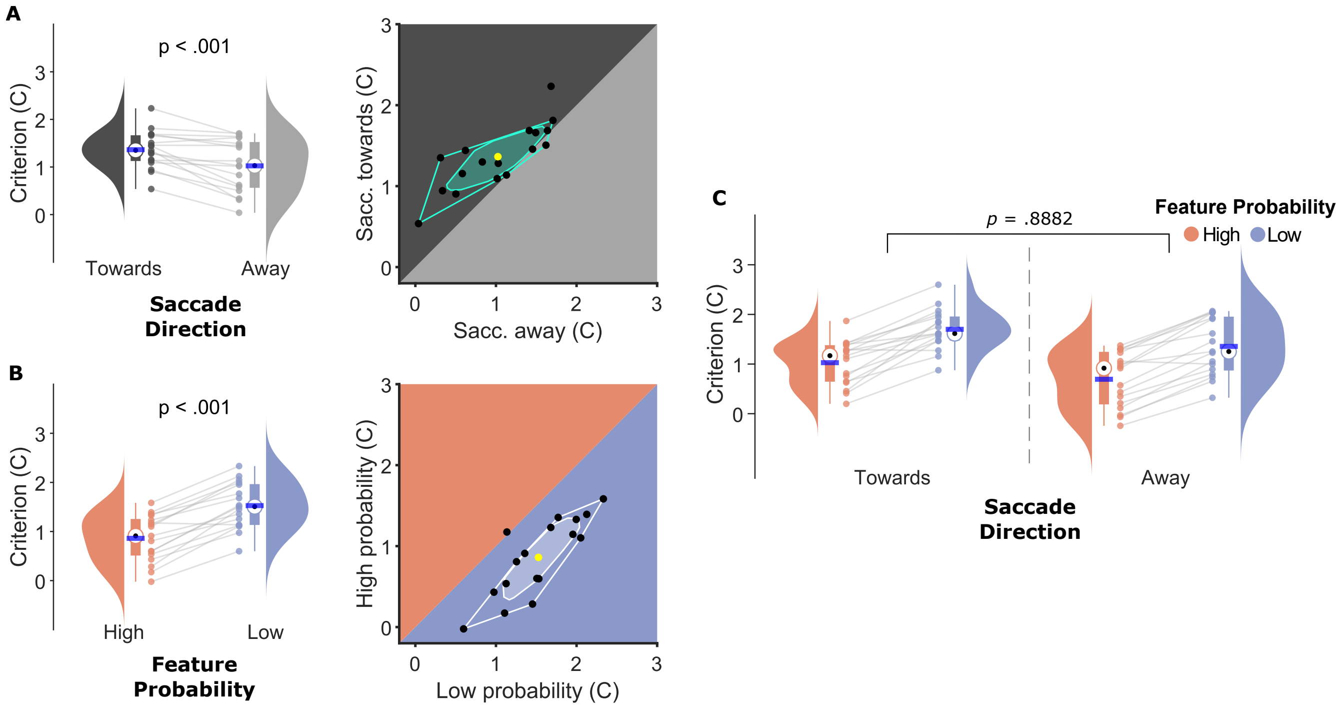


**Figure 1: Effects of presaccadic attention and expectation on Decision Criterion (C)**. **(A)** Criterion (C) as a function of saccade direction relative to target position: Towards and Away conditions are shown in dark gray and light gray, respectively. *Left panel:* Half-violin plots depict the distribution of criterion values across participants, with individual participants shown as dots. Boxplots show the interquartile range (IQR), whiskers extend to the most extreme values within 1.5 times the IQR, black dots indicate the median, and horizontal blue lines the mean. *Right panel:* Bagplot for Criterion ($C$) for saccades toward vs away from the response cue. The green polygon (bag) contains 50% of the data (black dots) whereas the surrounding fence, also in green, contains the remaining non-outlier data. The mean is indicated by the yellow dot. **(B)** Criterion (C) as a function of target probability: Target orientation with High and Low probability is shown in orange and blue, respectively. *Left panel:* Plotting conventions are the same as in A. *Right panel:* Bagplot for Criterion ($C$) for High and Low probability conditions. Parameters are the same as in A. Overall, less expected targets led to more conservative responses. **(C)** Criterion ($C$) as a function of target probability for each saccade direction: High and Low probability are shown in light red and blue, respectively.

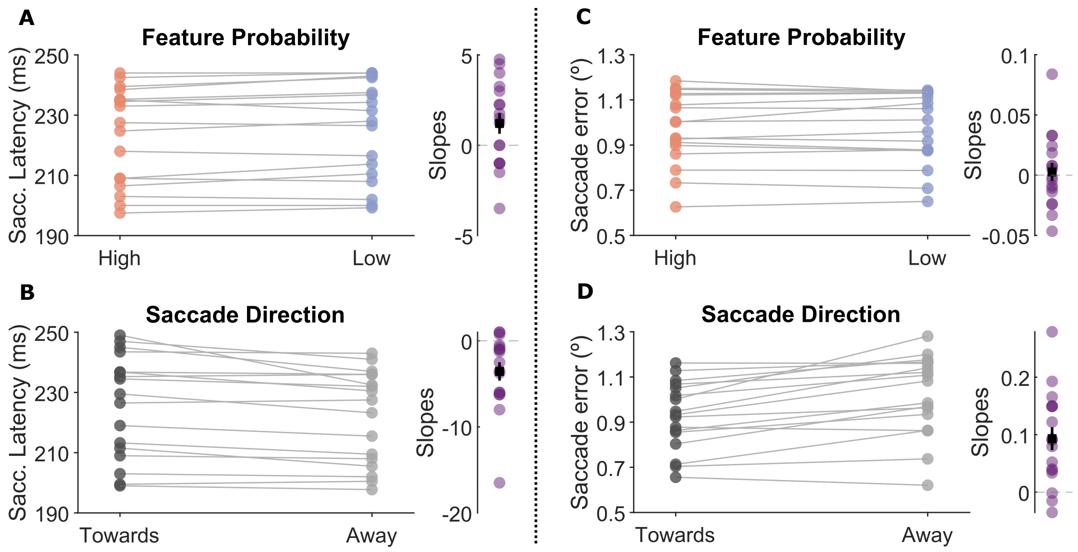


**Figure 2: Slopes derived from Feature Probability and Saccade Direction comparisons were used as covariates in separate ANCOVAs.**. **(A)** Saccade Latency (ms) as a function of feature probability (High and Low) and **(B)** Saccade Direction (Towards and Away). Each line connects each subject's mean between conditions. Regression slopes for each subject are shown in purple on the right, with black squares indicating the group mean slope and vertical black lines indicating the standard error (SE). Slopes from target probability and saccade direction were used as two separate covariates in the ANCOVA. **(C)** Saccade error (°) as a function of Feature Probability (High and Low) and **(D)** Saccade Direction (Towards and Away). The remaining parameters are the same as in A and B.

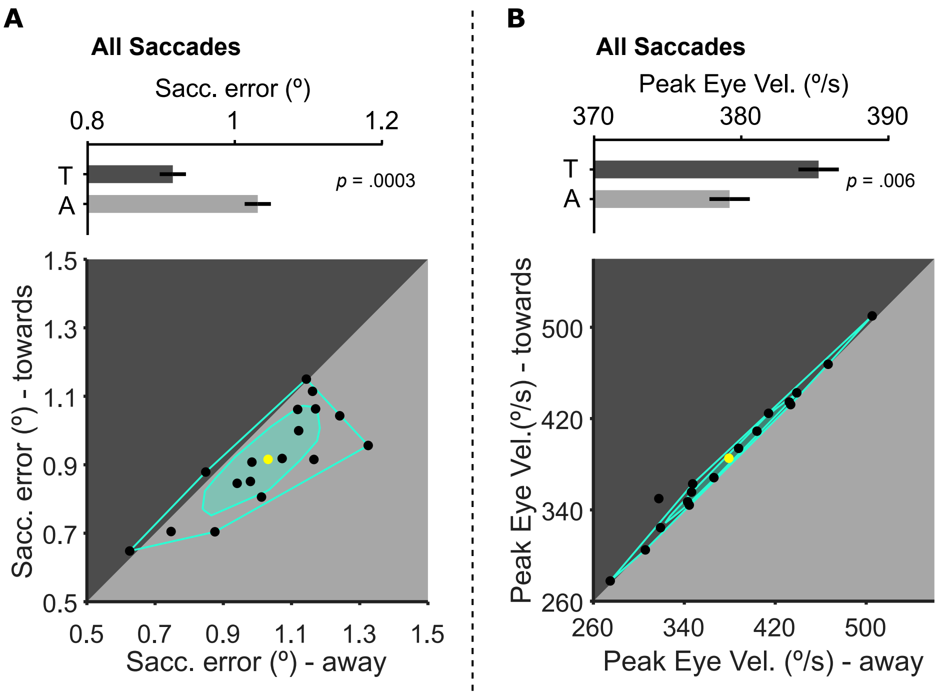


**Figure 3: Saccade endpoint error (°) and Peak eye Velocity (°/s) for signal-present trials**. Upper panels show mean (A) Saccade endpoint error (°) and (B) Peak eye Velocity (°/s), respectively, and lower panels show the corresponding bagplots using the same data. Error bars in the upper panels indicate the standard error of the mean (SEM). In the lower panels, black dots indicate individual participants, the polygon (bag) contains 50% of the observations, the surrounding fence contains the remaining non-outlier observations, and the yellow dot indicates the mean. The diagonal line indicates equal values between the two conditions. **(A)** Saccade endpoint error (°) and **(B)** Peak eye Velocity (°/s) as a function of saccade direction relative to the visual target location. Towards (T) and Away (A) conditions are shown in dark gray and light gray, respectively.

.3778).

*Within Subjects Effects*

| Cases | Sum of Squares | df | Mean Square | F | p | $\eta^2_p$ |
|---|---|---|---|---|---|---|
| Direction | 19.017 | 1 | 19.017 | 39.892 | < .001 | 0.714 |
| Residuals | 7.627 | 16 | 0.477 | | | |
| Probability | 12.850 | 1 | 12.850 | 28.453 | < .001 | 0.640 |
| Residuals | 7.226 | 16 | 0.452 | | | |
| Presaccadic Timing | 25.835 | 1 | 25.835 | 43.182 | < .001 | 0.730 |
| Residuals | 9.572 | 16 | 0.598 | | | |
| Direction ✻ Probability | 0.320 | 1 | 0.320 | 2.611 | .126 | 0.140 |
| Residuals | 1.963 | 16 | 0.123 | | | |
| Direction ✻ Presaccadic Timing | 6.300 | 1 | 6.300 | 42.812 | < .001 | 0.728 |
| Residuals | 2.355 | 16 | 0.147 | | | |
| Probability ✻ Presaccadic Timing | 0.741 | 1 | 0.741 | 5.025 | .040 | 0.239 |
| Residuals | 2.359 | 16 | 0.147 | | | |
| Direction ✻ Probability ✻ Presaccadic Timing | 0.005 | 1 | 0.005 | 0.030 | .865 | 0.002 |
| Residuals | 2.529 | 16 | 0.158 | | | |

*Note.* Type III Sum of Squares

**Table 1: Within-Subjects two-away repeated-measures ANOVA for visual sensitivity ($d'$).** The table reports the results for the factors Presaccadic Timing, with Early and Late conditions corresponding to targets presented farther from and closer to saccade onset, respectively; Saccade Direction (Towards and Away from the probed location) and Feature Probability (High and Low). $df$ = Degrees of Freedom; $F$ = F-values; $p$ = p-values; $\eta_p^2$ = partial eta squared.

*Post Hoc Comparisons - Direction ✻ Presaccadic Timing*

| | | Mean Difference | SE | df | t | Cohen's d | $p_{bonf}$ |
|---|---|---|---|---|---|---|---|
| Towards, Late | Away, Late | 1.178 | 0.157 | 16 | 7.502 | 1.276 | < .001 |
| | Towards, Early | 1.302 | 0.176 | 16 | 7.379 | 1.410 | < .001 |
| | Away, Early | 1.620 | 0.179 | 16 | 9.061 | 1.753 | < .001 |
| Away, Late | Towards, Early | 0.124 | 0.177 | 16 | 0.700 | 0.134 | 1.000 |
| | Away, Early | 0.441 | 0.113 | 16 | 3.914 | 0.478 | .007 |
| Towards, Early | Away, Early | 0.317 | 0.110 | 16 | 2.894 | 0.344 | .063 |

*Note.* P-value adjusted for comparing a family of 6 estimates.

*Note.* Results are averaged over the levels of: Probability

**Table 2: *Post hoc* comparisons between Presaccadic Timing (Early and Late condition) and Saccade Direction (Towards and Away) for visual sensitivity ($d'$).** $SE$ = Standard Error; $df$ = Degrees of Freedom; $t$ = t-values; $p_{\text{bonf}}$ = Bonferroni corrected p-value.

*Post Hoc Comparisons - Probability ✻ Presaccadic Timing*

| | | Mean Difference | SE | df | t | Cohen's d | $p_{bonf}$ |
|---|---|---|---|---|---|---|---|
| High, Late | Low, Late | -0.467 | 0.152 | 16 | -3.076 | -0.506 | .043 |
| | High, Early | 1.019 | 0.128 | 16 | 7.945 | 1.103 | < .001 |
| | Low, Early | 0.257 | 0.128 | 16 | 2.015 | 0.278 | .366 |
| Low, Late | High, Early | 1.486 | 0.213 | 16 | 6.969 | 1.609 | < .001 |
| | Low, Early | 0.724 | 0.166 | 16 | 4.374 | 0.784 | .003 |
| High, Early | Low, Early | -0.762 | 0.110 | 16 | -6.909 | -0.825 | < .001 |

*Note.* P-value adjusted for comparing a family of 6 estimates.

*Note.* Results are averaged over the levels of: Direction

**Table 3: *Post hoc* comparisons between Presaccadic Timing (Early and Late condition) and Feature Probability (High and Low) for visual sensitivity ($d'$).** $SE$ = Standard Error; $df$ = Degrees of Freedom; $t$ = t-values; $p_{\text{bonf}}$ = Bonferroni corrected p-value.

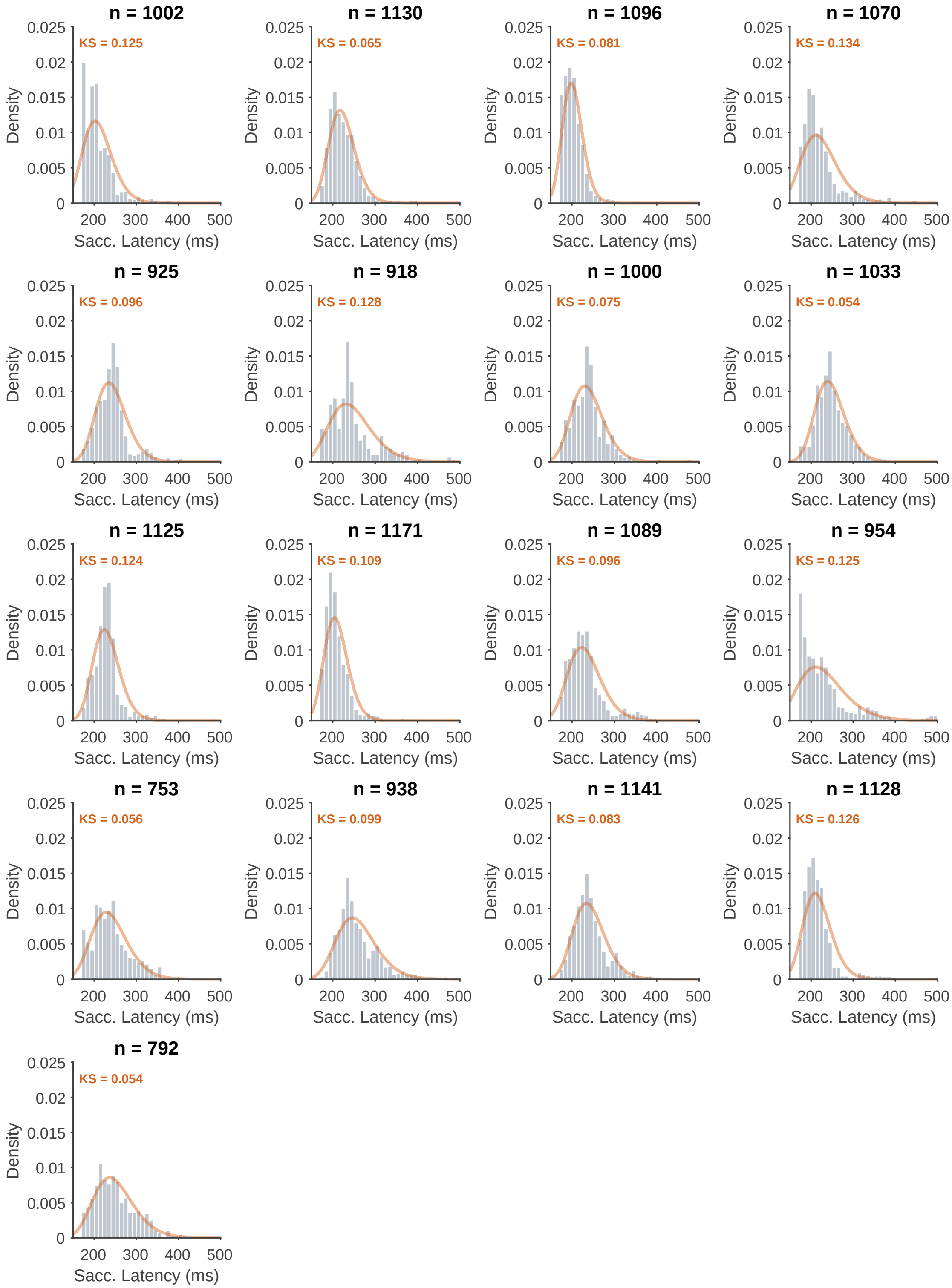


**Figure 4: Inverse-Gaussian fits to saccade-latency distributions.** Observed saccade-latency distributions are shown separately for each participant for signal trials only. Histograms show the empirical distribution of saccade latencies, and overlaid curves show the best-fitting inverse-Gaussian distribution fitted separately to each participant's latency distribution. The inverse-Gaussian distribution provided a reasonable approximation to the observed positively skewed latency distributions (Kolmogorov–Smirnov statistic: mean = $.096$, range = $.054$–$.134$), supporting its use as the response distribution in the generalized linear mixed model.
